# Utilising nuclear encoded plastid DNA to identify donors of grass-to-grass lateral gene transfer

**DOI:** 10.64898/2026.08.26.747220

**Authors:** Noah G. Bourne, Layla Payne, Sophie Manzi, Guillaume Besnard, Maria S. Vorontsova, Richard W. Jobson, Guillaume Chomicki, Luke T. Dunning

## Abstract

Determining the correct donor species/lineages of grass-to-grass lateral gene transfer (LGT) is vital for deducing specific donor features that could help inform the mechanism of transfer. This requires a dataset spanning a broad range of species to achieve the phylogenetic resolution necessary for precise donor inference. As grass-to-grass LGT often involves the transfer of multi-gene DNA fragments, they can contain additional sequences that allow for accurate orthologous comparisons, such as nuclear DNA of plastid origin (NUPTs). Here we systematically scan for NUPTs in the genomes of four *Alloteropsis semialata* accessions, whose LGTs have previously been characterised. Using the abundant Panicoideae chloroplast sequences, we reconstruct NUPT phylogenies and infer two lateral acquisitions: one from Paniceae/*Digitaria* and another from Andropogoneae/*Eremochloa* adjacent to a previously identified LGT. We then assembled and included an additional 12 *Eremochloa* chloroplast genomes in the analysis and showed the likely donor was *Eremochloa attenuata*. Subsequent short-read mapping from *E. attenuata* to the nuclear region flanking this NUPT showed consistent coverage across the region, including the previously identified LGT, supporting co-transfer. Overall this study highlights the potential for NUPTs to better identify the donors of grass-to-grass LGT.

## Introduction

Lateral gene transfer (LGT) is the transfer of DNA between two species outside of sexual reproduction, and its inference typically relies on phylogenetic analyses. Early phylogenetic studies were usually based on DNA sequences from organelle genomes (Palmer & Zamir, 1982; Brown et al., 1982). These organelle genomes were also among the first to be fully sequenced (Anderson et al., 1981; Shinozaki et al., 1986). This in turn meant that early studies in which the first cases of eukaryotic LGTs were identified initially characterised multiple accounts of endosymbiotic gene transfer (Gray, 1992). Subsequent studies showed that eukaryotic LGTs extended far beyond just organelle endosymbiotic gene transfer s (Andersson, 2005) but was still largely limited to microbial eukaryotes (Keeling and Palmer, 2008, Etten and Bhattacharya, 2020). As more eukaryotic genomes have become available, the prevalence and evolutionary significance of eukaryotic LGTs has become apparent and acts on a much broader scale, involving all major eukaryotic lineages (Li *et al*., 2022, Widen *et al*., 2023, Keeling, 2024, Policarpo *et al*., 2025, Mariault *et al*., 2025). Despite this, key limitations remain. For example, identifying the exact donor can be challenging because donors may be extinct, particularly for older transfer events, or may have subsequently lost the native transferred gene themselves. Moreover, even when potential extant donors can be identified, confirming their role is often limited by the availability of sequence data, both for the putative donor and for closely related species. This is because accurate inference requires strong support for the transferred gene nesting within the donor clade.

Accurate donor assignment is essential for downstream analyses, particularly when identifying shared genomic and phenotypic features among donor lineages to better understand the mechanisms that facilitate these transfers. Phylogenetic methods traditionally rely on coding portions of single genes to infer LGT events, which requires extensive sampling of that homologue. In grass-to-grass LGT however, it has been well established that DNA can be transferred via large multigene fragments which often include non-coding intronic and intergenic DNA (Dunning *et al*., 2019, Hibdige *et al*., 2021, Mahelka *et al*., 2021, Raimondeau *et al*., 2023). Therefore there is scope to identify additional DNA sequence features within the fragments that have been more widely sampled, such as rDNA (Mahelka *et al*., 2017) and Nuclear DNA of Plastid Origin (NUPT).

Endosymbiotic gene transfer between chloroplasts and their concurrent nuclear genomes has been a continuous phenomenon since the primary endosymbiosis of cyanobacteria into an ancient eukaryote (Timmis *et al*., 2004, Bock, 2017). While many of these early transfers led to substantial gene loss within the proto-organelle genomes, as their functions were effectively outsourced to the nucleus (Zhong, Kuijl-van den Berg and Chung, 2026), it was generally thought most chloroplast insertions would be neutral or slightly deleterious. More recently however, NUPTs have been shown to play functional roles through the transfer of both coding and non-coding sequences (Carretero-Paulet, Marczuk-Rojas and Gálvez-Salido, 2026). NUPTs of various sizes have been identified in almost all sequenced plant genomes to date, ranging from under one hundred base pairs to the insertion of entire chloroplast genomes (Huang *et al*., 2017, Marczuk-Rojas, Maldonado and Carretero-Paulet, 2025). Although NUPTs make up an overall small proportion of nuclear genome content, approximately 0.02-3.29% of angiosperm genomes (Marczuk-Rojas, Maldonado and Carretero-Paulet, 2025), their continued acquisition and conservation (due to functional co-option) make them an everpresent feature of plant genomes. Grass chloroplast genomes have been far more extensively assembled than their nuclear counterparts. As of April 2026, a search of the NCBI Assembly database using the query ‘Poaceae AND chloroplast AND complete genome’ returned records for 1343 species compared to ∼136 nuclear genomes sampled in 2024 (Pereira *et al*., 2026). While there is a seemingly low chance that NUPTs could be incorporated into LGT fragments, species involved in multiple lateral transfer events increases the likelihood of co-transfer. NUPTs that were inserted into the donor genome prior to transfer are ideal candidates for phylogenetic inference of LGTs for the following five reasons: 1) There is a much larger dataset in which to infer potential donors to the LGT fragments; 2) Grass chloroplasts are generally conserved (Hu *et al*., 2022), which means the vast majority of NUPTs will have homologous sequences across grasses; 3) While they vary in size, many NUPTs contain ample sequence information for meaningful phylogenetic inference; 4) Chloroplast-derived sequences are less affected by paralogy and genome duplication than nuclear genes (pre-transfer), simplifying evolutionary inference; and 5) NUPTs are relatively easy to identify with a simple BLAST search.

This study utilises four previously assembled *A. semialata* genomes that have been extensively screened for LGTs, with transferred fragments already characterised (Raimondeau *et al*., 2023). To see if NUPTs can be reliably used to better identify LGT donors, they were first identified in the *A. semialata* genomes and then we reconstructed their evolutionary histories using phylogenetic analyses to assess potential lateral acquisition. Subsequently, the location of laterally acquired NUPTs is examined to determine whether they occur near or within previously identified LGT fragments, or represent potential novel fragments. Finally, the inferred donor lineages of the nuclear LGTs are compared with those of their corresponding NUPTs to identify a potential common origin.

## Methods

### NUPT identification pipeline

BLASTN v2.15.0 (Camacho *et al*., 2009) databases were built for each of the four *Alloteropsis semialata* nuclear genomes from (Raimondeau *et al*., 2023). The *A. semialata* chloroplast along with 395 other Panicoideae chloroplast genomes were then BLAST against each assembly reporting hits with a minimum e-value of 1 x 10^-10^. Panicoideae was chosen as the vast majority of nuclear LGTs (92.2%) were acquired from this subfamily (Raimondeau *et al*., 2023). To ensure there would be sufficient information for accurate phylogenetic analysis the BLAST hits were filtered to include matches that were at least 1 kb and a minimum of 95% percentage identity. A custom python script then assessed chloroplast–nuclear hits to compare whether ‘foreign’ (non-*A. semialata*) chloroplast sequences produced stronger matches to the same nuclear regions than the *A. semialata* chloroplast. This initial Panicoideae dataset excluded other *Alloteropsis* species to prevent ‘foreign’ hits from sister or closely related taxa, as LGT between *Alloteropsis* species cannot be reliably inferred at this phylogenetic resolution. BLAST results (outfmt 6) from self (*A. semialata*) and foreign chloroplast queries were parsed and grouped by nuclear subject sequence. For each self hit, overlapping foreign hits (minimum 50-bp overlap) to the same nuclear region were identified. The best foreign hit was then selected amongst all foreign hits based on highest bit score, followed by percentage identity as a secondary criterion. Following this, the top foreign hit was considered to outperform the self hit only if its bit score exceeded the self hit by at least 100. This threshold represents a substantial difference in alignment significance and was chosen to ensure that only strong candidates are retained. Results were recorded per nuclear region, including alignment statistics and produced an output file that inferred candidate NUPTs of potential foreign origin (**Supplementary Table S1**). Finally, the FASTA sequences of the foreign NUPTs were then extracted from their respective genomes using BEDtools v2.3.1 (Quinlan and Hall, 2010) and taken forward for phylogenetic analysis (**Figure 1A**).

**Figure 1.**
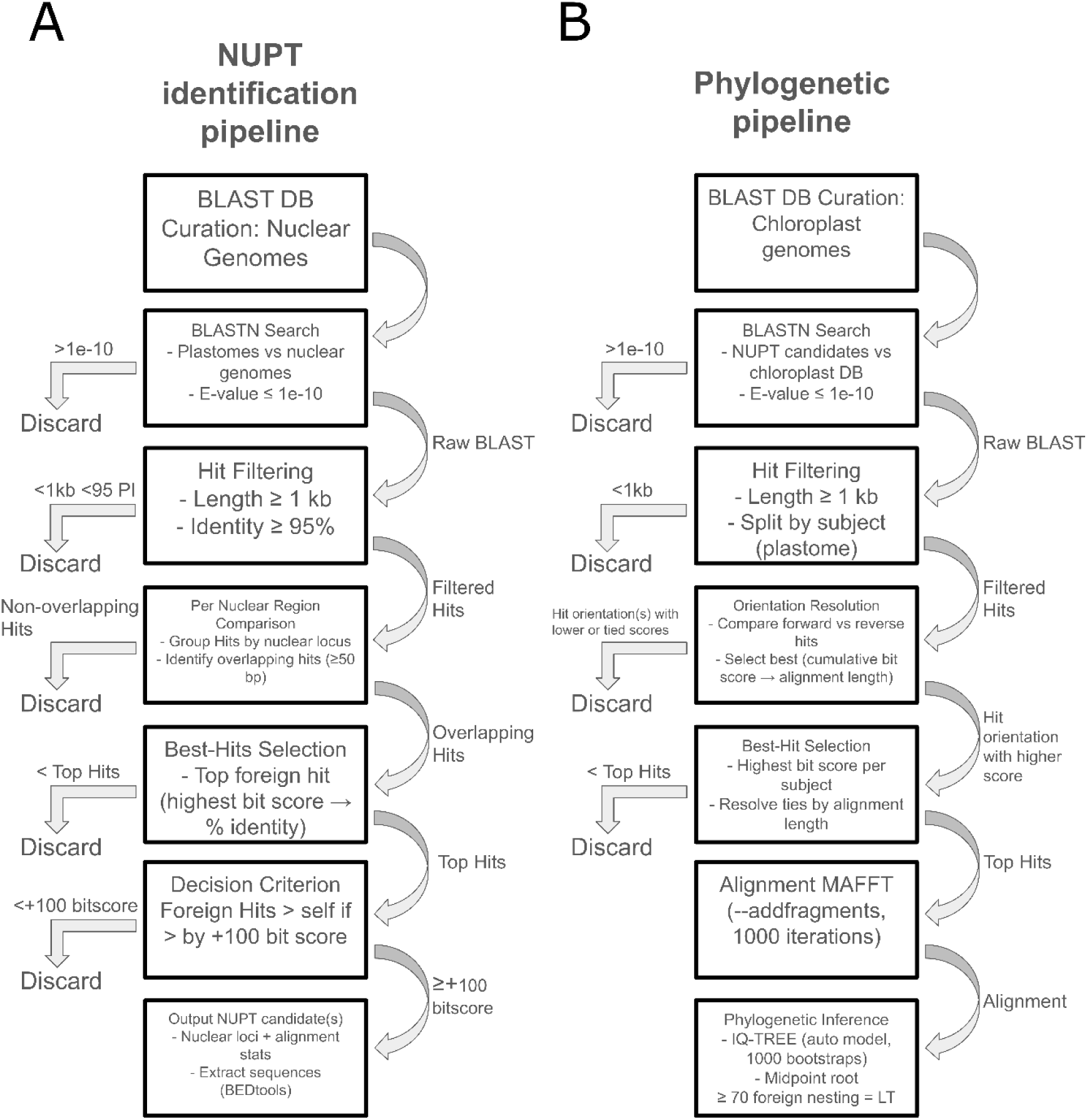
Flowcharts of the NUPT identification and phylogenetic pipelines. PI = percentage identity, DB = Database, LT = Lateral transfer. The output of **A)** ‘NUPT identification pipeline’ is parsed directly into **B)** ‘Phylogenetic pipeline’. Alt text: Step by step flowchart of how the NUPT identification and Phylogenetic pipelines were executed.

### Chloroplast genome sequencing and assembly

Genomic DNA of 12 *Eremochloa* herbarium samples (**Supplementary Table S2**) was extracted with the BioSprint DNA Plant Kit (Qiagen Inc.). Libraries were constructed using the TruSeq Nano DNA Sample Preparation Kit (Illumina, San Diego, CA, USA) as detailed in Bianconi *et al*. (2020) using 50 to 500 ng of double-stranded DNA per sample. Sequencing was performed at the Get-PlaGe core facility, Castanet-Tolosan, France, on the Illumina HiSeq 3000 platform, generating 150-bp paired-end reads through bridge amplification.

Chloroplast genomes were assembled from the short reads of the *Eremochloa* accessions using the program Getorganelle v1.7.7.1 (Jin *et al*., 2020). Initial assemblies were generated using default *k*-mer sizes (21, 45, 65, 85 and 105), word sizes (estimated), and extending contigs for 15 rounds with *E. ophiuroides* (NC_036716.1), *E. ciliaris* (NC_035028.1) and *E. eriopoda* (NC_035023.1) chloroplast assemblies as reference seeds. If the initial assemblies were under 120 kb and or fragmented the parameters were adjusted according to recommendations from https://github.com/Kinggerm/GetOrganelle/wiki/FAQ, and then subsequently re-assembled until a single circular contig or linear scaffold(s) >120kb (if complete circular assemblies were unattainable) was produced. The chloroplast genomes were then annotated on GeSeq (Tillich *et al*., 2017) web server using the same three *Eremochloa* chloroplasts as reference.

### NUPT phylogenetic pipeline

The 12 newly assembled *Eremochloa* chloroplast genomes were then combined with 402 other chloroplast genomes (downloaded from NCBI) from across Panicoideae, this included the chloroplast genomes from 395 Panicoideae accessions plus the four available *Alloteropsis* accessions (*A. semialata*, *A. cimicina, A. angusta*, and *A. paniculata*) and three Chloridoideae outgroups (**Supplementary Table S3**). A BLAST database of all 414 chloroplast genomes was first constructed. The NUPT candidates were then BLAST against this dataset again using a minimum e-value of 1 x 10^-10^. BLAST outputs were then split by subjects and filtered, only retaining hits of a minimum of 1000 bp. Due to the presence of the inverted repeat regions in chloroplast genomes, many returned similar or identical BLAST matches in multiple orientations. To account for this, forward and reverse strand hits were compared within accessions based on cumulative bitscore and then, in the case of a tie, total alignment length of all BLAST hits representing that orientation. If orientations could not be separated by these metrics they were not taken forward in the analysis to prevent comparisons with orientations that may represent non-orthologous regions. In cases where the best supported orientation was selected, the single top BLAST hit (if multiple hits for that orientation) with the highest bitscore was taken forward again using alignment lengths in the cases of ties. The raw hits per subject were also output in separate files for downstream manual checks to ensure the correct hit was selected. The resulting hits from each NUPT query were then aligned using MAFFT v7.453 (Katoh and Standley, 2013) using the –addfragments parameter and 1000 iterations. Finally, phylogenetic trees were inferred for each alignment using IQ-Tree v2.3.0 (Minh *et al*., 2020) with automatic model selection and 1000 bootstrap replicates. The NUPT phylogenies were then midpoint rooted and manually inspected on Figtree v1.4.4 (http://tree.bio.ed.ac.uk/software/figtree/). A NUPT was considered laterally acquired if it nested within other well supported groups outside of its species topology and with a bootstrap score ≥70 (**Figure 1B, Supplementary Tables S4**). Laterally transferred NUPTs identified in the initial phylogenetic analysis were subsequently re-examined by inspecting the hit subject coordinates on the putative donor chloroplast genome. If hits were found to derive from contiguous regions of the chloroplast genome, the corresponding continuous NUPT regions were concatenated and the phylogenetic analysis was repeated. Alignments in these cases were trimmed on Geneious to remove regions with >50% gaps to ensure accurate branch lengths. If the contiguous NUPT phylogenies remained consistent with lateral acquisition they were then manually re-rooted at the species within Chloridoideae.

### Short read mapping

Bowtie2 v2.5.4 (Langmead and Salzberg, 2012) was used to index the AUS1 *A. semialata* genome, and short reads were aligned using default parameters. Regions spanning 200 kb upstream and downstream of the LGT locus (ASEM_AUS1_21433) were extracted from BAM files and converted to BigWig format using deepTools v3.5.6 (Ramírez et al., 2014). Coverage tracks were normalized to 1× genome coverage using the reads per genomic content (RPGC) method with a bin size of 10 bp. The resulting BigWig files were imported into R v4.4.2 (R core team, 2024) and visualized using the ggcoverage package (Song and Wang, 2023).

## Results

### Thousands of NUPTs were identified in the Alloteropsis semialata genomes

The initial NUPT scan for each of the four *A. semialata* nuclear genomes yielded between 1084 and 1420 hits when its own chloroplast genome was BLAST against the assemblies. We then repeated the BLAST search with 398 additional chloroplast genomes from across Panicoideae. These were then filtered and potential ‘foreign’ NUPTs were inferred based on whether there were stronger matches to chloroplast genomes outside of *A. semialata* (see *Methods*). Overall, between 6 and 56 foreign NUPT candidates were identified per accession. Of these, only four displayed phylogenetic topologies consistent with lateral acquisition (**Supplementary File S1**), with two identified in each of the AUS1 and TAN1-04b accessions (**Table 1**).

**Table 1.** Statistics from the NUPT identification and phylogenetic pipeline.

| Accession | Total self hits | Filtered NUPTs | Foreign NUPTs | Phylogenetically verified NUPTs |
| --- | --- | --- | --- | --- |
| AUS1 | 1285 | 27 | 6 | 2 |
| RSA5-3 | 1228 | 233 | 56 | 0 |
| TAN1-04B | 1420 | 172 | 53 | 2 |
| ZAM1505-10 | 1084 | 129 | 12 | 0 |

To inspect the chloroplast regions in which the NUPTs showing topologies consistent with lateral acquisition were derived, the four NUPTs were subsequently BLAST against their respective hypothesised donor chloroplast genomes. In both AUS1 and TAN1-04b genomes, their two NUPTs were from contiguous regions (**Figure 2**). The concatenated NUPTs in TAN1-04B return a 3919-bp BLAST match (E-value = 0, 98% percentage identity) to a contiguous region within the large single copy (LSC) region of *Digitaria sanguinalis.* The AUS1 concatenated NUPTs return a 3348-bp BLAST match (E-value = 0, 96.4% percentage identity) in the small single copy (SSC) region of *Eremochloa ciliaris*.

**Figure 2.**
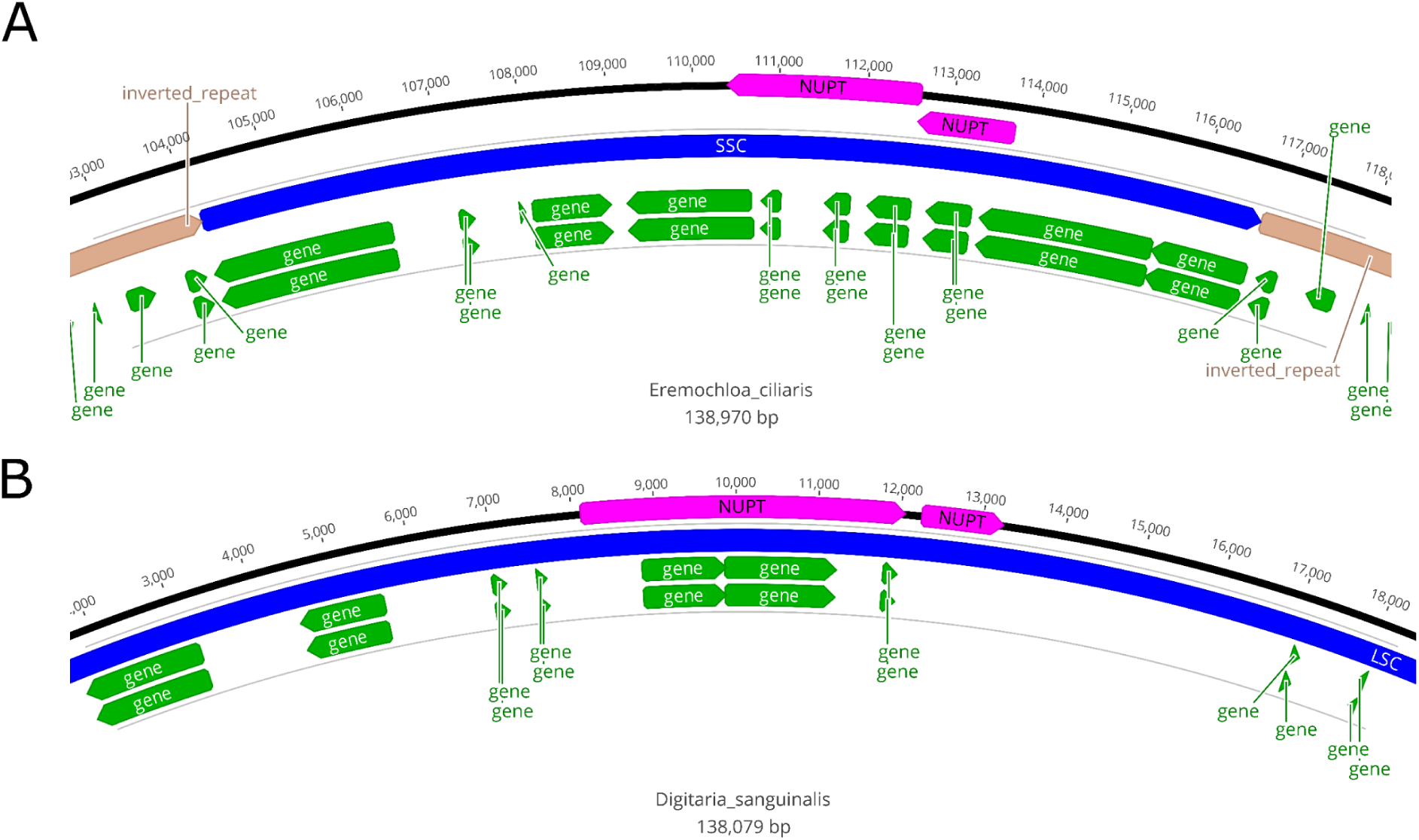
Schematic showing BLAST matches between concatenated contiguous NUPTs of *Alloteropsis semialata* and chloroplast regions of a putative donor. The inner green arrows represent chloroplast genes. The outer blue arrows represent small single copy (SSC) (**A**) and large single copy (LSC) regions (**B**). Finally, the light brown arrows represent distinct inverted repeat chloroplast regions. The pink arrows represent the respective BLAST matches from *A. semialata* NUPTs. **A)** AUS1 nested NUPTs mapped to the *Eremochloa ciliaris* chloroplast genome. **B)** TAN1-04B nested NUPTs mapped to the *Digitaria sanguinalis* chloroplast genome. Alt text: Diagram showing the chloroplast regions, in each donor species, from which the laterally transferred NUPTs originated.

### Multiple NUPTs have been laterally transferred into Alloteropsis semialata

The phylogenetic pipeline was re-run on the respective concatenated NUPTs from AUS1 and TAN1-04B. The TAN1-04B concatenated NUPT continued to show high supported placement outside of the species topology with it again being sister to Anthephorinae*/Digitaria* species within Paniceae (**Figure 3**). Further evidence that this was a laterally acquired NUPT is found in the placement of the *A. semialata* native chloroplast region along with the other *Alloteropsis* accessions, which follow the expected species topology, notably nesting in Paniceae/Boivinellinae (**Figure 3**). The TAN1-04B NUPT was found on a 96-kb scaffold and was found to overlap with two annotated gene models (TAN1-04B_38564 & TAN1-04B_38565). Interestingly these gene models or the surrounding genes do not appear in Raimondeau *et al*. (2023) analysis as LGTs, so this NUPT likely represents the remnants of a potentially unidentified LGT fragment in the genome.

**Figure 3.**
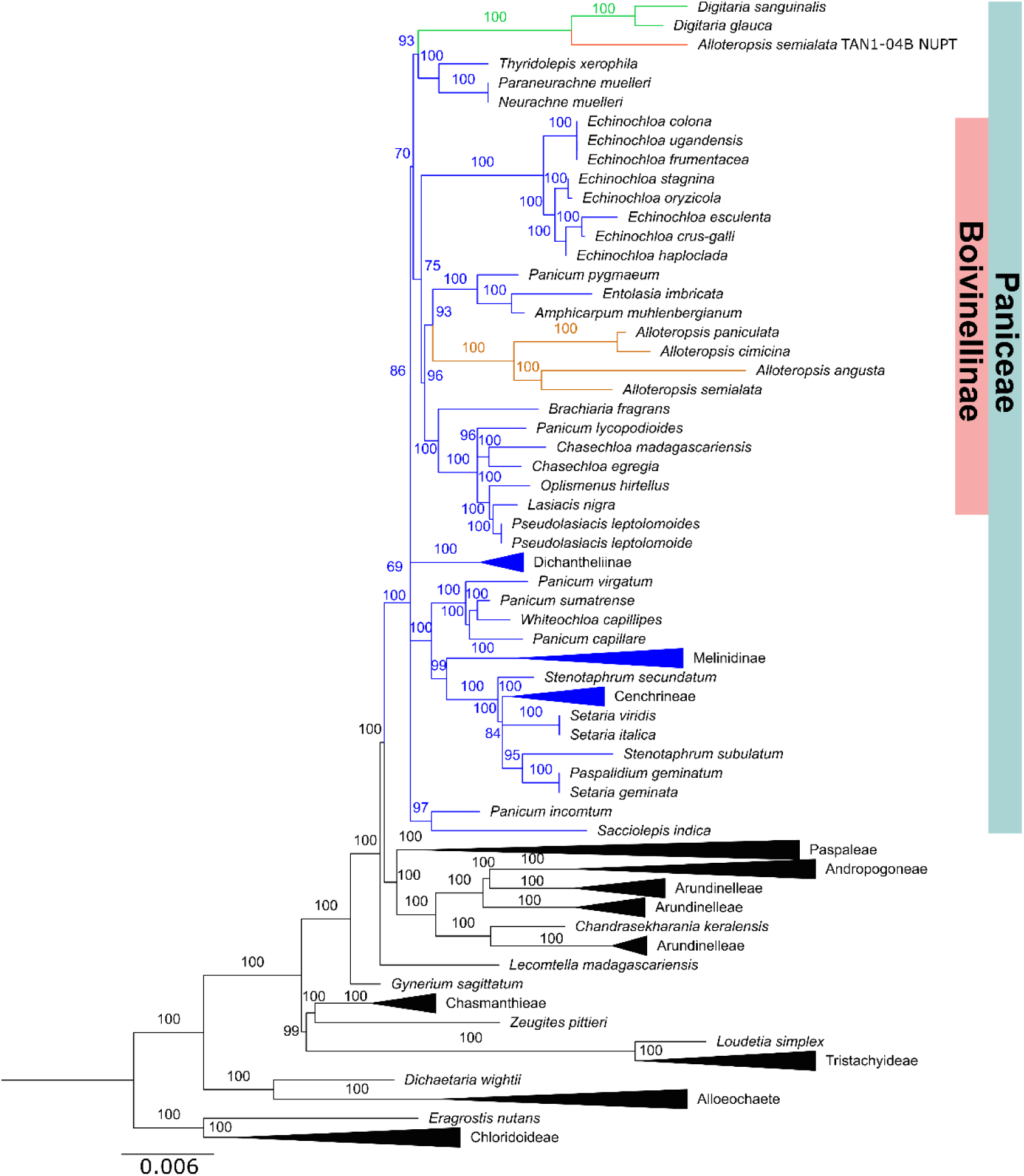
Maximum likelihood phylogeny of all homologous chloroplast regions to the combined TAN1-04B NUPT. The substitution model selected was TPM2u+F+I+R4 (see *Methods*). The clade/branches are highlighted in orange for natives (e.g. *A. semialata* expected species topology), green for donors and red for the NUPT. The Paniceae tribe branches are in blue. Grey branches represent species falling outside of their expected species topology most likely due to the fact the tree is built from an alignment of only 3930 bp, so some incongruence can be expected. The scale bar is set to branch lengths. The phylogeny shows the highly supported placement of the TAN1-04B NUPT sister to *Digitaria* species outside of its expected species topology which would be nesting in Boivinellinae. Alt text: Phylogenetic tree showing a NUPT identified in the *Alloteropsis semialata* TAN1-04B accession does not nest within the other *Alloteropsis* species and is more closely related to *Digitaria* species. The phylogenetic tree has good support, suggesting this region was laterally transferred and not vertically inherited.

The AUS1 initial NUPT trees suggested that an Andropogoneae/Ratzeburgiinae/*Eremochloa* species was the likely donor as both NUPTs were sister to the clade. To more accurately infer the donor, 12 additional *Eremochloa* chloroplast genomes were assembled from paired-end short read data. Eleven chloroplast genomes represented complete circular assemblies with only one accession, *Eremochloa lanceolata*, assembling a single non-circular scaffold. Despite this, the scaffold was still of comparable size to the other complete genomes and only annotated one less gene (**Table 2**).

**Table 2.** Assembly size and gene count for newly assembled *Eremochloa* chloroplast genomes.

| *Eremochloa* species | Complete/scaffold | Contigs | Genome size | Gene count |
| --- | --- | --- | --- | --- |
| *E. attenuata* | Complete | 1 | 139,399 | 138 |
| *E. bimaculata* | Complete | 1 | 138,881 | 138 |
|  | Complete | 1 | 139,003 | 138 |
|  | Complete | 1 | 138,988 | 138 |
| *E. ciliaris* | Complete | 1 | 138,874 | 138 |
| *E. eriopoda* | Complete | 1 | 138,887 | 138 |
| *E. lanceolata* | Scaffold | 1 | 138,828 | 137 |
| *E. muricata* | Complete | 1 | 139,386 | 138 |
| *E. ophiuroides* | Complete | 1 | 139,129 | 138 |
|  | Complete | 1 | 139,107 | 138 |
| *E. petelotii* | Complete | 1 | 140,307 | 138 |
| *E. zeylanica* | Complete | 1 | 139,342 | 138 |

Taking the AUS1 concatenated NUPT and the additional *Eremochloa* chloroplast assemblies, the phylogenetic pipeline was re-run. The resulting tree displayed the AUS1 concatenated NUPT nesting with high support within the *Eremochloa* clade (bootstrap = 100) (**Figure 4**). The NUPT was placed sister to *E. attenuata* (bootstrap = 100) indicating that this species is or a very close relative of the NUPT donor. This lateral transfer event was also supported by the native *A. semialata* chloroplast region nesting with the other *Alloteropsis* species in their species topology (**Figure 4**).

**Figure 4.**
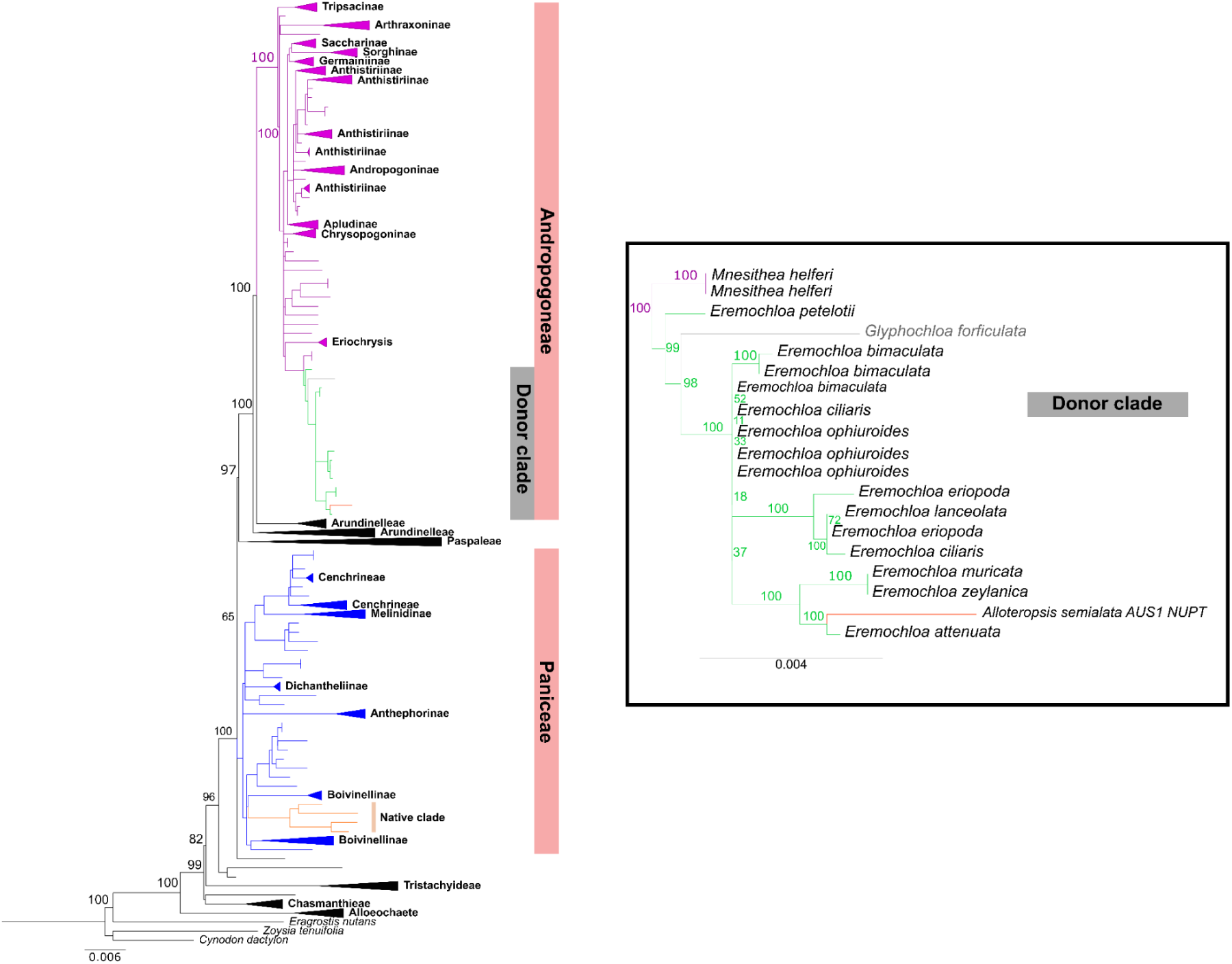
Maximum likelihood phylogeny of all homologous chloroplast regions to the combined AUS1 NUPT. The substitution model selected was TVM+F+I+R4 (see *Methods*). The clade/branches are highlighted in orange for natives (e.g. *A. semialata* expected species topology), green for donors and red for the NUPT. The Paniceae and Andropogoneae tribe branches are in blue and purple, respectively. Grey branches represent species falling outside of their expected species topology most likely due to the fact the tree is built from an alignment of only 3264 bp, so some incongruence can be expected. The scale bar is set to branch lengths. The black box highlights the zoomed-in section of the donor clade (Andropogoneae/Ratzeburgiinae) and laterally transferred NUPT. Alt text: Phylogenetic tree showing a NUPT identified in the *Alloteropsis semialata* AUS1 accession does not nest within the other *Alloteropsis* species but with *Eremochloa* species. The phylogenetic tree has good support, suggesting this region was laterally transferred and not vertically inherited.

Following the phylogenetic analysis, supporting that the AUS1 concatenated NUPT was laterally acquired, its location was checked relative to other LGTs in the AUS1 genome assembly. The NUPTs were then found to be within the vicinity (∼65 kb downstream) of a previously identified LGT (ASEM_AUS1_21433) from Raimondeau *et al*. (2023). The donor of this nuclear gene was inferred to be in Andropogoneae, which is consistent with the NUPT phylogeny. To see if the NUPT and nuclear LGT could have been transferred as one fragment and thus potentially had a common origin, *Eremochloa attenuata* short reads were mapped to the region surrounding the nuclear LGT up until the closest laterally transferred NUPT (**Figure 5**). The *E. attenuata* short reads mapped consistently across the initially identified LGT and into the surrounding genes and intergenic DNA. This suggests this region along with the genes is likely present in *E. attenuata* and could have been transferred together as one would not usually expect relatively consistent homology to both genes and surrounding intergenic DNA. The NUPT was characterised by high coverage mapping across multiple species in the plot reflecting the often high coverage nature of chloroplast DNA even when sequencing low coverage. Interestingly the other species in which we mapped as controls also mapped over the LGT and two downstream genes indicating that they could be present in these species as well.

**Figure 5.**
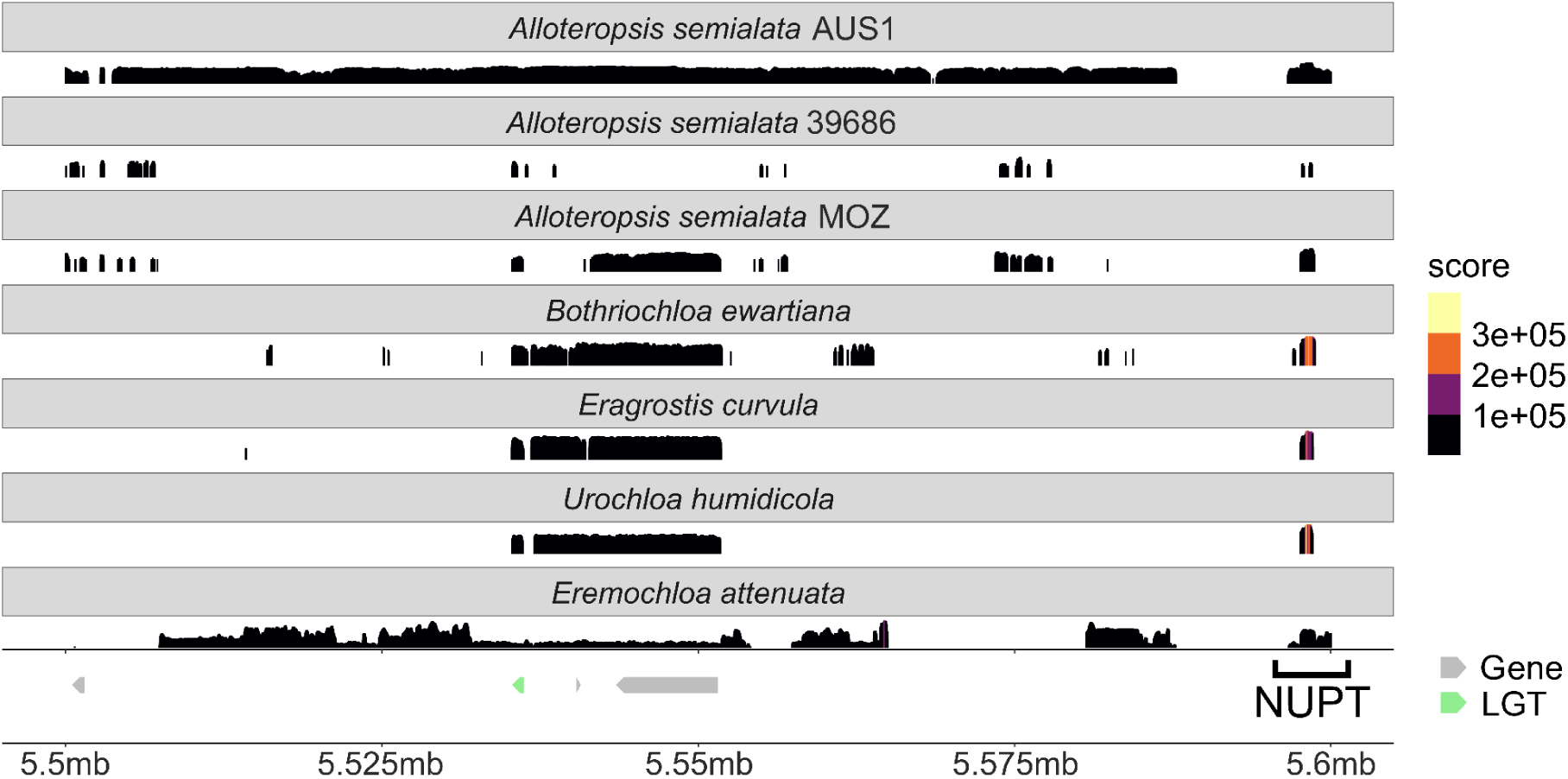
Short read mapping to the AUS1 LGT region. Short read mapping to the region surrounding the LGT ASEM_AUS1_21433 from fragment AUS1_FRAG_05 identified on chromosome 3 in Raimondeau *et al*. (2023). The initial LGT is highlighted in green with one of the corresponding NUPTs discovered in this study labelled downstream. The score has been calculated from 1× RPCG normalisation (see *Methods*) and represents the relative enrichment or depletion of reads at that position. Alt text: Graphical representation of a zoomed in 100 kilobase region that encompasses the AUS1 laterally transferred NUPT and previously identified laterally transferred nuclear gene. The black histogram in each lane represents short reads mapped from those respective species to the region. The figure is designed to show *Eremochloa attenuata’s* consistent mapping to the originally identified LGT and surrounding region, providing evidence alongside the phylogenetic analysis, that this region could represent a laterally transferred DNA fragment.

## Discussion

Donor inference of grass-to-grass LGT through phylogenetic analysis of nuclear genes is often constrained by the limited availability of nuclear genome assemblies. Here, we demonstrate the use of NUPTs as an additional approach for donor identification due to the greater availability of chloroplast sequence data enabling a more accurate inference of the donor lineage. Secondly, the study highlights the ease of incorporating additional data, as assembling chloroplast genomes requires only low-coverage whole-genome sequencing (WGS) and the high conservation grass chloroplast genomes means that it is very likely to assemble the region of interest. While it is not certain that *Eremochloa attenuata* is the true donor, as the transfer could have been from a yet to be sequenced *Eremochloa* species and/or ancestor, the resolution of donor lineage identification was still significantly improved. Moreover through read mapping, we show how donor inference from the NUPT can be extended to nearby genes that have or have not been previously identified as LGT. While phylogenetic evidence is still required to confirm common origin in the adjacent nuclear genes, the read mapping at least provides candidate genes/regions that could have arrived as part of the same fragment. Interestingly, the read mapping suggests that this specific LGT fragment may have been independently transferred multiple times into distantly related grass lineages, similar to the C_4_ genes described by Christin et al. (2012). This possibility warrants further investigation but is beyond the scope of this work.

NUPTs are readily identifiable within the genome, providing an opportunity to detect previously unrecognised laterally transferred DNA beyond nuclear genes. This includes the use of non-coding regions for phylogenetic inference, as NUPT insertions often span multiple genes, including introns and intergenic sequences. Because NUPTs are integrated as discrete fragments, the full sequence they contain can be used for reliable orthologous comparisons. For example, the AUS1 and TAN1-04B NUPTs encompass multiple genes as well as non-coding DNA and successfully generated highly supported phylogenies. Importantly this region was ignored in Raimondeau *et al*. (2023) as these chloroplast genes were not initially annotated as gene models in the nuclear assemblies, so were not present in the cds FASTA files in which the previous LGT inference methods were based on. Beyond previously identified LGT fragments, the TAN1-04B NUPT highlighted a potential new LGT fragment that has been previously uncharacterised due to a lack of nearby nuclear genes showing a clear signal of lateral transfer. This now opens up the potential for analysis of laterally transferred DNA remnants that may not include nuclear genes or are largely degenerated.

It is important to acknowledge that NUPTs are highly variable between plants (Marczuk-Rojas, Maldonado and Carretero-Paulet, 2025) so the likelihood of a NUPT being transferred within a DNA fragment may be extremely low in some species. Furthermore, if the LGT mechanism does not involve DNA intermediates, for example, the lateral transfer of mRNA (Yoshida *et al*., 2010), then again the utility of NUPTs, in the context of transferring alongside nuclear genes, is negligible (unless a transcribed NUPT itself is transferred). Finally, this method is likely only sensitive enough for recent transfers, as NUPTs are likely to degenerate unless they are of functional significance in the recipients genome (Carretero-Paulet, Marczuk-Rojas and Gálvez-Salido, 2026) with estimates from *Oryza sativa subsp. japonica* showing ∼80% of NUPTs degrade within a million years of integration (Matsuo *et al*., 2005).

The NUPT identification and phylogenetic pipeline could be easily extended beyond chloroplast sequences and incorporate nuclear DNA of mitochondrial origin (NUMTs) which are also prevalent across plants (Zhang *et al*., 2020). However, due to the complexity of many plant mitochondrial genomes (Gualberto and Newton, 2017) the available sequences are not as extensive as chloroplast sequences. Although new tools are now able to consistently assemble plant mitogenomes (Zhou *et al*., 2025) meaning their availability will likely substantially increase. The use of NUMTs to infer lateral acquisition would in theory be applicable to a much larger set of organisms outside of grasses and even plants. Of course it is unclear whether other LGT mechanisms outside of grasses involve the transfer of large DNA fragments that would incorporate NUMTs, however, the methodology, if such a case was identified, would remain consistent.

The results here also point to the potential of DNA to traverse large evolutionary distances through non-vertical, multistep processes. The case presented in this work includes, organelle-to-nuclear transfer followed by nuclear-to-nuclear transfer. Additionally, there is an example of non-vertical multistep transfers in parasitic plants, involving the intracellular gene transfer of a plastid sequence to the mitochondrial genome in a *Cuscuta* species, which was subsequently laterally transferred again, through a host intermediate, to another parasitic plant, *Orobanche rigens* (Denysenko-Bennett et al., 2025). Both of these examples demonstrate that the lateral acquisition of DNA is not constrained to a single transfer event which should be taken into consideration especially when inferring donor lineages.

Overall this study provides evidence for the effective incorporation of NUPTs into phylogenetic pipelines to help identify the donors of grass-to-grass LGT. The pipeline in this study displays the potential for the generation of high supported phylogenies that span a wide range of species due to the large database of chloroplast sequences. Finally, this study shows scope for the incorporation of NUMTs into future analysis and continues to underscore the importance of organelle genomes for phylogenetic inference.

## Acknowledgements

We gratefully acknowledge the curators and staff of the Queensland Herbarium (BRI) and Royal Botanic Gardens, Kew Herbarium (K) for providing access to their collections and specimens for this study.

## Author contributions

Conceptualization: N.B and L.D. Methodology: N.B. Data curation: N.B., R.W.J., M.S.V., S.M., G.B and L.D. Data analysis: N.B and L.P. Interpretation of data: N.B. Figures: N.B. Writing: N.B and L.D with the help of all authors.

## Data availability statement

The raw reads and chloroplast assemblies from the *Eremochloa* accessions have been uploaded to NCBI under the bioproject number PRJNA1492527. Multiple sequence alignments and corresponding phylogenetic trees have been uploaded to Dryad under DOI: 10.5061/dryad.v15dv42bz. The scripts can be found at https://github.com/NBourne04/Identification-and-phylogenetic-analysis-of-NUPTs.git

## Supplementary Data

**Supplementary Table S1:** BLAST comparison results (see Methods: NUPT identification pipeline) for NUPTs per accession.

**Supplementary Table S2:** Details of Eremochloa species sample collection

**Supplementary Table S3:** Chloroplast accessions used in the study

**Supplementary Table S4:** Results from the NUPT phylogenetic pipeline per accession

**Supplementary Table S5:** Details of accessions used for short read mapping

**Supplementary Files S1:** Alignments and phylogenetic trees used to generate the NUPT phylogenies per accession. Dryad Reviewer Link: http://datadryad.org/share/LINK_NOT_FOR_PUBLICATION/_HQbYCh6EjiJQNMxeG5SVVGO0V0z3KCBowCVGmxXFWw

## References

Anderson, S., Bankier, A. T., Barrell, B. G. et al. (1981) ‘Sequence and organization of the human mitochondrial genome’, Nature, 290(5806), pp. 457–465. Available at: 10.1038/290457a0.

Andersson, J. O. (2005) ‘Lateral gene transfer in eukaryotes’, Cellular and Molecular Life Sciences, 62(11), pp. 1182–1197. Available at: 10.1007/s00018-005-4539-z.

Bianconi, M. E., Hackel, J., Vorontsova, M. S. et al. (2020) ‘Continued adaptation of C4 photosynthesis after an initial burst of changes in the Andropogoneae grasses’, Systematic Biology, 69(3), pp. 445–461. Available at: 10.1093/sysbio/syz066.

Bock, R. (2017) ‘Witnessing genome evolution: experimental reconstruction of endosymbiotic and horizontal gene transfer’, Annual Review of Genetics, 51, pp. 1–22. Available at: 10.1146/annurev-genet-120215-035329.

Brown, W.M., Prager, E.M., Wang, A. et al. (1982) ‘Mitochondrial DNA sequences of primates: Tempo and mode of evolution’, Journal of Molecular Evolution 18, 225–239. 10.1007/BF01734101

Camacho, C., Coulouris, G., Avagyan, V. et al. (2009) ‘BLAST+: architecture and applications’, BMC Bioinformatics, 10(1), p. 421. Available at: 10.1186/1471-2105-10-421.

Carretero-Paulet, L., Marczuk-Rojas, J. P. and Gálvez-Salido, A. (2026) ‘Nuclear DNA of plastid origin (NUPTs), neglected driver of genome variation and evolutionary innovation’, The Plant Journal, 125(3), p. e70685. Available at: 10.1111/tpj.70685.

Christin, P.-., Edwards, E. J., Besnard, G. et al. (2012) ‘Adaptive Evolution of C4 Photosynthesis through Recurrent Lateral Gene Transfer’, Current Biology, 22(5), pp. 445–449. Available at: 10.1016/j.cub.2012.01.054.

Denysenko-Bennett, M., Kwolek, D., Góralski, G. et al. (2025) ‘Horizontal gene transfer of the Pytheas sequence from Cuscuta to Orobanche via a host-mediated pathway’, Scientific Reports, 16(1), p. 2056. Available at: 10.1038/s41598-025-31853-x.

Dunning, L. T., Olofsson, J. K., Parisod, C. et al. (2019) ‘Lateral transfers of large DNA fragments spread functional genes among grasses’, Proceedings of the National Academy of Sciences of the United States of America, 116(10), pp. 4416–4425. Available at: 10.1073/pnas.1810031116.

Etten, J. V. and Bhattacharya, D. (2020) ‘Horizontal gene transfer in Eukaryotes: not if, but how much?’, Trends in Genetics, 36(12), pp. 915–925. Available at: 10.1016/j.tig.2020.08.006.

Gray, M. W. (1992) ‘The endosymbiont hypothesis revisited’, International Review of Cytology. Academic Press, pp. 233–357. Available at: 10.1016/S0074-7696(08)62068-9.

Gualberto, J. M. and Newton, K. J. (2017) ‘Plant mitochondrial genomes: dynamics and mechanisms of mutation’, Annual Review of Plant Biology, 68, pp. 225–252. Available at: 10.1146/annurev-arplant-043015-112232.

Hibdige, S. G. S., Raimondeau, P., Christin, P.-. et al. (2021) ‘Widespread lateral gene transfer among grasses’, New Phytologist, 230(6), pp. 2474–2486. Available at: 10.1111/nph.17328.

Hu, Y., Sun, Y., Zhu, Q.-. et al. (2022) ‘Poaceae chloroplast gnome sequencing: great leap forward in recent ten years’, Current Genomics, 23(6), pp. 369–384. Available at: 10.2174/1389202924666221201140603.

Huang, Y., Wang, J., Yang, Y. et al. (2017) ‘Phylogenomic analysis and dynamic evolution of chloroplast genomes in Salicaceae’, Frontiers in Plant Science, 8, 1050. Available at: 10.3389/fpls.2017.01050.

Jin, J.-., Yu, W.-., Yang, J.-. et al. (2020) ‘GetOrganelle: a fast and versatile toolkit for accurate de novo assembly of organelle genomes’, Genome Biology, 21, p. 241. Available at: 10.1186/s13059-020-02154-5.

Katoh, K. and Standley, D. M. (2013) ‘MAFFT Multiple Sequence Alignment Software Version 7: Improvements in Performance and Usability’, Molecular Biology and Evolution, 30(4), pp. 772–780. Available at: 10.1093/molbev/mst010.

Keeling, P. J. and Palmer, J. D. (2008) ‘Horizontal gene transfer in eukaryotic evolution’, Nature Reviews Genetics, 9(8), pp. 605–618. Available at: 10.1038/nrg2386.

Keeling, P. J. (2024) ‘Horizontal gene transfer in eukaryotes: aligning theory with data’, Nature Reviews Genetics, 25(6), pp. 416–430. Available at: 10.1038/s41576-023-00688-5.

Langmead, B. and Salzberg, S. L. (2012) ‘Fast gapped-read alignment with Bowtie 2’, Nature Methods, 9(4), pp. 357–359. Available at: 10.1038/nmeth.1923.

Li, Y., Liu, Z., Liu, C. et al. (2022) ‘HGT is widespread in insects and contributes to male courtship in lepidopterans’, Cell, 185(16), pp. 2975–2987.e10. Available at: 10.1016/j.cell.2022.06.014.

Mahelka, V., Krak, K., Kopecký, D. et al. (2017) ‘Multiple horizontal transfers of nuclear ribosomal genes between phylogenetically distinct grass lineages’, Proceedings of the National Academy of Sciences, 114(7), pp. 1726–1731. Available at: 10.1073/pnas.1613375114.

Mahelka, V., Krak, K., Fehrer, J. et al. (2021) ‘A Panicum-derived chromosomal segment captured by Hordeum a few million years ago preserves a set of stress-related genes’, The Plant Journal, 105(5), pp. 1141–1164. Available at: 10.1111/tpj.15167.

Marczuk-Rojas, J. P., Maldonado, A. D. and Carretero-Paulet, L. (2025) ‘Episodic and ongoing mechanisms drive plastid-derived nuclear DNA evolution in Angiosperms’, Genome Biology and Evolution, 17(11), p. evaf194. Available at: 10.1093/gbe/evaf194.

Mariault, L., Puginier, C., Keller, J. et al. (2025) ‘Mechanisms, detection, and impact of horizontal gene transfer in plant functional evolution’, The Plant Cell, 37(9), p. koaf195. Available at: 10.1093/plcell/koaf195.

Matsuo, M., Ito, Y., Yamauchi., R., et al. (2005) ‘The Rice Nuclear Genome Continuously Integrates, Shuffles, and Eliminates the Chloroplast Genome to Cause Chloroplast–Nuclear DNA Flux’, The Plant Cell, 17(3), pp. 665–675. Available at: 10.1105/tpc.104.027706.

Minh, B. Q., Schmidt, H. A., Chernomor, O. et al. (2020) ‘IQ-TREE 2: New Models and Efficient Methods for Phylogenetic Inference in the Genomic Era’, Molecular Biology and Evolution, 37(5), pp. 1530–1534. Available at: 10.1093/molbev/msaa015.

Palmer, J.D. & Zamir, D. (1982) ‘Chloroplast DNA evolution and phylogenetic relationships in Lycopersicon’, Proceedings of the National Academy of Sciences of the United States of America, 79(16), pp. 5006–5010, 10.1073/pnas.79.16.5006.

Pereira, L., Alenazi, A. S., Mian, S. et al. (2026) ‘Gene turnover in the common ancestor of all C4 grasses’, Plants, People, Planet [in press]. Available at: 10.1002/ppp3.70206.

Policarpo, M., Salzburger, W., Maumus, F. et al. (2025) ‘Multiple horizontal transfers of immune genes between distantly related teleost fishes’, Molecular Biology and Evolution, 42(5), p. msaf107. Available at: 10.1093/molbev/msaf107.

Quinlan, A.R. and Hall, I.M. (2010) ‘BEDTools: a flexible suite of utilities for comparing genomic features’, Bioinformatics, 26(6), pp. 841–842. Available at: 10.1093/bioinformatics/btq033.

R Core Team (2024). _R: A Language and Environment for Statistical Computing. R Foundation for Statistical Computing, Vienna, Austria. <https://www.R-project.org/>.

Shinozaki, K., Ohme, M., Tanaka, M. et al. (1986) ‘The complete nucleotide sequence of the tobacco chloroplast genome: its gene organization and expression’, The EMBO Journal, 5(9), pp. 2043–2049. Available at: 10.1002/j.1460-2075.1986.tb04464.x.

Song, Y. and Wang, J. (2023) ‘ggcoverage: an R package to visualize and annotate genome coverage for various NGS data’, BMC Bioinformatics, 24(1), p. 309. Available at: 10.1186/s12859-023-05438-2.

Raimondeau, P., Bianconi, M. E., Pereira, L. et al. (2023) ‘Lateral gene transfer generates accessory genes that accumulate at different rates within a grass lineage’, New Phytologist, 240(5), pp. 2072–2084. Available at: 10.1111/nph.19272.

Ramírez, F., Dündar, F., Diehl, S. et al. (2014) ‘deepTools: a flexible platform for exploring deep-sequencing data’, Nucleic Acids Research, 42(W1), pp. W187–W191. Available at: 10.1093/nar/gku365.

Tillich, M., Lehwark, P., Pellizzer, T. et al. (2017) ‘GeSeq – versatile and accurate annotation of organelle genomes’, Nucleic Acids Research, 45(W1), pp. W6–W11. Available at: 10.1093/nar/gkx391.

Timmis, J. N., Ayliffe, M. A., Huang, C. Y. et al. (2004) ‘Endosymbiotic gene transfer: organelle genomes forge eukaryotic chromosomes’, Nature Reviews Genetics, 5(2), pp. 123–135. Available at: 10.1038/nrg1271.

Widen, S. A., Bes, I. C., Koreshova, A. et al. (2023) ‘Virus-like transposons cross the species barrier and drive the evolution of genetic incompatibilities’, Science, 380(6652), p. eade0705. Available at: 10.1126/science.ade0705.

Yoshida, S., Maruyama, S., Nozaki, H. et al. (2010) ‘Horizontal Gene Transfer by the Parasitic Plant Striga hermonthica’, Science, 328(5982), pp. 1128–1128. Available at: 10.1126/science.1187145.

Zhang, G.-., Dong, R., Lan, L.-. et al. (2020) ‘Nuclear integrants of organellar DNA contribute to genome structure and evolution in plants’, International Journal of Molecular Sciences, 21(3), p. 707. Available at: 10.3390/ijms21030707.

Zhong, Y., Kuijl-van den Berg, F. and Chung, K. P. (2026) ‘Plastid genes in motion: mechanisms and physiological implications of endosymbiotic gene transfer’, Journal of Experimental Botany, 77(10), pp. 2810–2814. Available at: 10.1093/jxb/erag086.

Zhou, C., Brown, M., Blaxter, M. et al. (2025) ‘Oatk: a de novo assembly tool for complex plant organelle genomes’, Genome Biology, 26(1), p. 235. Available at: 10.1186/s13059-025-03676-6.

